# Optimization of isolation, expansion, and differentiation of canine intestinal organoids

**DOI:** 10.64898/2026.03.19.712113

**Authors:** Inês E. Dias, Arabela Ritchie, Maarten Delemarre, Kerstin Schneeberger, Carlos A. Viegas, Isabel R. Dias, Pedro P. Carvalho, Bart Spee

## Abstract

Intestinal organoids are three-dimensional *in vitro* structures derived from stem cells and serve as a valuable model for studying intestinal biology and pathophysiology. This study optimized the isolation, expansion, and differentiation of canine intestinal organoids from duodenum and colon. Organoids were generated from canine intestinal crypts and cultured in Matrigel with a growth factor cocktail. The impact of prostaglandin E_2_ (PGE_2_) concentration on organoid growth was evaluated, and a two-phase differentiation protocol—comprising patterning and differentiation media—was implemented, including interleukin (IL)-22 in the duodenal differentiation phase. Organoids cultured with 100 nM PGE_2_ exhibited increased crypt budding and organoid-forming efficiency, indicative of enhanced stem cell proliferation. Differentiated organoids expressed key intestinal markers (VIL1, SI, CHGA, MUC2), and forskolin-induced swelling demonstrated functional Cystic Fibrosis Transmembrane Conductance Regulator (CFTR) activity. Although the sample size (n=2) limits generalizability, this optimized protocol provides a relevant *in vitro* model for studying canine intestinal function. The model can be used in future research for disease modelling and translational applications, supporting downstream studies in gastrointestinal disease, drug permeability, and comparative One Health research.

## 1. Introduction

Organoid technology has revolutionized research on developmental biology, disease modelling, and therapeutic applications across various species, including humans and animals (1,2). Intestinal organoids, have emerged as powerful *in vitro* models that closely mimic the architecture and functionality of the intestinal epithelium. These 3D structures, derived from stem cells, provide a unique opportunity to study the physiology and pathology of the small and large intestine (3–5). While organoid models of murine intestine have been extensively studied and characterized, canine intestinal organoids remain comparatively underexplored (6). They hold substantial promise for translational research, especially given the natural occurrence of gastrointestinal diseases in dogs, such as inflammatory bowel disease (IBD) and colorectal cancer (7,8). These conditions share clinical and pathological features with human counterparts, making canine organoid models particularly valuable for understanding disease mechanisms and evaluating potential therapeutic interventions, with less ethical concerns. Among the canine intestinal segments, the duodenum and colon are of particular interest due to their distinct roles in nutrient absorption, immune modulation, and disease processes (7,9).

Significant advances in organoid research have introduced methods for the long-term culture of isolated intestinal crypts or stem cells. When provided with an optimized growth factor cocktail (comprising epidermal growth factor (EGF), Noggin, and R-spondin-1) and grown in a three-dimensional extracellular matrix such as Matrigel, these stem cells can develop into organoids that mimic the native intestinal epithelium (7,8,10). These “mini-guts” exhibit essential functions of the intestinal tissue and serve as robust models for investigating molecular regulatory and pathological mechanisms of the intestinal epithelium (7,11–13). Pioneering work by Sato *et al*. (2009) demonstrated that single LGR5+ cells isolated from murine intestinal crypts could form genetically stable, self-renewing 3D structures *in vitro* (11). These organoids replicated key features of the intestinal crypt-villus architecture and differentiated into various cell types, including enterocytes, goblet cells, Paneth cells, and enteroendocrine cells (11,14). This culture approach has since been extended to multiple species, including humans, rats, dogs and cats (3,8,15–18). Figure 1 illustrates the various cell types present in the canine intestine (duodenum and colon). This schematic represents a generalized organization of the canine intestinal epithelium and reflects conserved epithelial lineages described across mammalian species.

**Figure 1.**
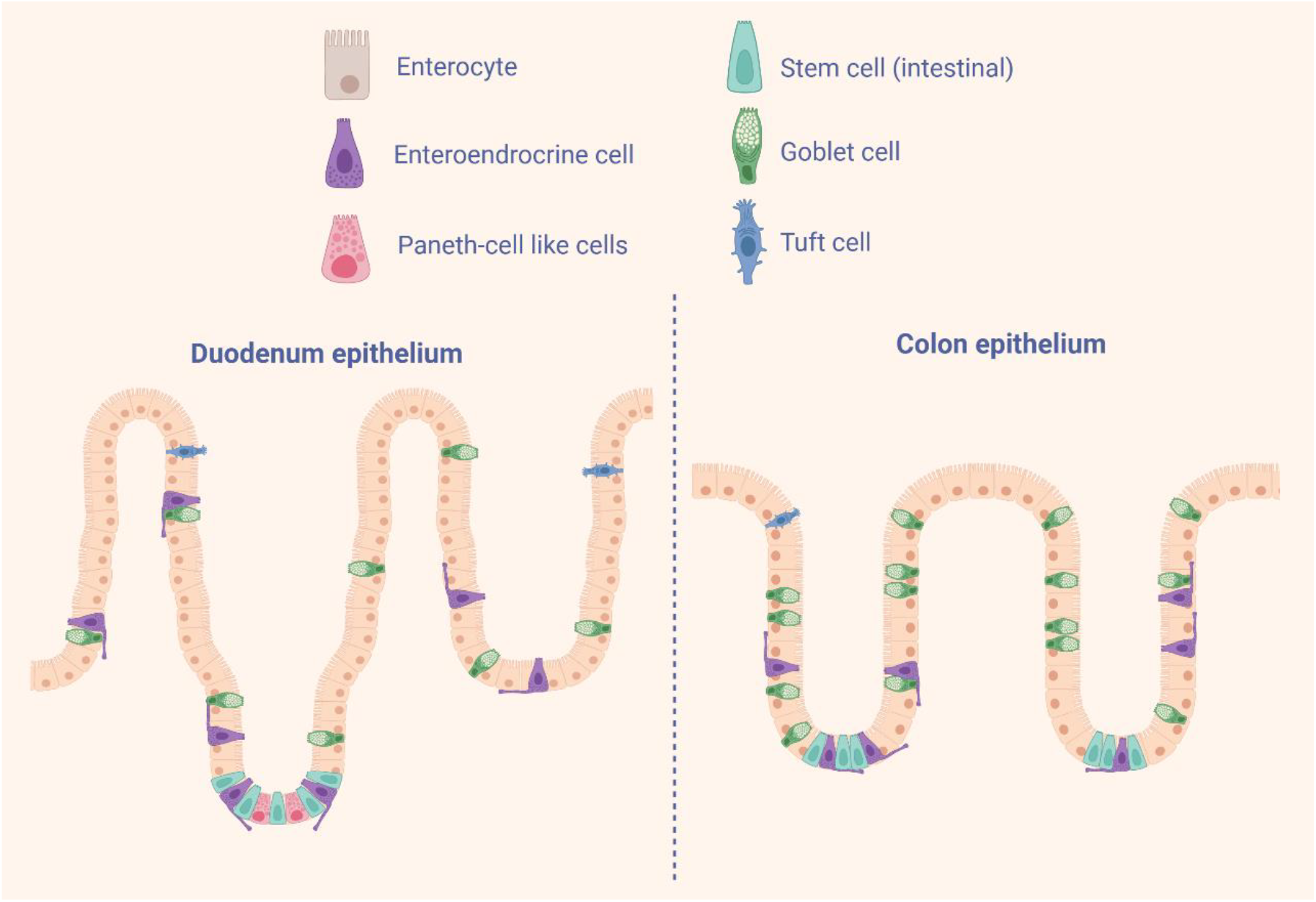
Overview of the major epithelial cell types present in the canine intestinal epithelium. Schematic illustration of the duodenal and colonic epithelia showing the distribution of enterocytes, goblet cells, enteroendocrine cells, Paneth-like cells, tuft cells, and intestinal stem cells. (Original figure created with BioRender.com).

Despite their potential, there is a significant lack of research on the isolation, expansion, and differentiation of canine intestinal organoids. While some studies have reported successful culture and basic characterization of these organoids (15,19,20), evidence regarding their differentiation into specialized intestinal cell types remains limited and poorly defined. Standard expansion media tend to favor stem cell maintenance at the expense of secretory lineage differentiation, whereas differentiation-promoting conditions may reduce viability (19). In addition, the cellular mechanisms supporting the intestinal stem cell niche in dogs remain incompletely defined. Chandra *et al*. (2019) reported absence of classical Paneth cell markers such as lysozyme, while detecting expression of crypt-associated markers including EPHB2 and FZD5, suggesting that niche composition in dogs may differ from that described in murine and human systems (15). Recent advancements in human organoids have linked IL-22 to enhance epithelial regeneration and promote Paneth cell-associated functions (21). IL-22 primarily supports epithelial defense by promoting Paneth cell formation and inducing antimicrobial gene expression across multiple intestinal epithelial cell types, without directly driving stem cell expansion or absorptive lineage differentiation (6,22,23). He *et al*. (2022) also incorporated PGE_2_ into the expansion medium (EM) for organoid culture (21). Previous studies have demonstrated that PGE_2_ directly stimulates epithelial cells, leading to robust activation of ion transport and fluid secretion, thereby regulating epithelial function independently of immune cells (24). Combined, this study optimized culture conditions to enhance stem cell proliferation and promote differentiation into specialized cell types. To achieve this, they developed a two-step differentiation strategy, beginning with a patterning medium (PM) to transition the cells from a proliferative state to early differentiation, mimicking the intestinal crypt niche environment. This was followed by a differentiation medium (DM) to promote lineage-specific maturation, resulting in cellular compositions that closely resemble the native intestinal epithelium (21). In dogs, there is currently no evidence confirming the presence of Paneth cells in the intestine (15,25). However, studies suggest the existence of Paneth-like cells or other epithelial cell types capable of producing antimicrobial peptides, a function typically associated with Paneth cells in humans and murine models (15,25). Based on these observations, we explored whether supplementation with IL-22 and PGE_2_ could improve proliferation and differentiation efficiency in canine duodenal and colonic organoids.

This study aims to address existing gaps by optimizing methods for the isolation, expansion, and differentiation of canine intestinal organoids, focusing on duodenal and colonic tissues. The novel expansion and differentiation media were tested on canine organoids, and a two-phase differentiation protocol, consisting of PM followed by DM, was applied to the organoid models, as previously established in human studies (21). We hypothesize that applying human-derived culture and differentiation protocols to canine intestinal tissues will enhance stem cell proliferation and support the generation of functionally and morphologically mature organoids. By advancing the characterization and differentiation protocols of canine intestinal organoids, this research aims to develop reliable models of the canine intestinal epithelium. These foundational advancements will enable future studies, including the development of disease models and comparative analyses with human intestinal diseases.

## 2. Methods

### 2.1. Animals

Duodenal and colonic samples were obtained as surplus material from two fresh canine cadavers used in unrelated research, which was performed according to the Dutch Experiments on Animals Act and conformed to EU standards (European Directive 2010/63/EU). The selected dog had no history of gastrointestinal issues, and mortality was associated with non-intestinal factors. One animal was male and the other female. The samples were collected less than 12h post-mortem.

### 2.2. Isolation, Cultivation and Cryopreservation of Canine Intestinal Organoids

The isolation method was adapted from Van der Hee *et al*. (2020) (13). A 2 cm section of the duodenum and colon was dissected and immersed in ice-cold PBS. After opening the sections longitudinally, they were washed three times with ice-cold Phosphate buffered saline (PBS) supplemented with 1% (v/v) penicillin-streptomycin and 0.25 µg/ml amphotericin-B (Gibco, Life Technologies). The mucosa was separated from the muscular layer using a surgical knife, then cut into small cubes. To ensure thorough cleaning, the fragments were placed in a 15 mL Falcon tube containing Dulbecco’s modified Eagle’s medium (DMEM) F12 supplemented with 10% (v/v) fetal bovine serum (FBS) and 1% (v/v) penicillin-streptomycin and 0.25 µg/ml amphotericin-B, and subjected to repeated gentle aspiration and expulsion using a pipette. The cleaned fragments were then transferred to ice-cold PBS containing 30 mM EDTA and incubated for 30 minutes with gentle rotation at room temperature. Following this, vigorous pipetting was performed (approximately 40 repetitions) to dislodge the crypts using a 1,000 µl pipette tip. The suspension was centrifuged at 500 g for 5 minutes at 4°C, and the supernatant was discarded. The crypt pellet was resuspended in Matrigel and plated into pre-warmed 24-well plates. Empty 24-well suspension plates had been pre-incubated at 37°C overnight in a CO_2_ incubator to enhance surface tension.

Organoids were passaged at a 1:4 ratio every 7 days and culture medium was refreshed every 2-3 days. Organoids were passaged by enzymatic digestion, by collecting the organoids in ice-cold DMEM, followed by centrifugation at 500 g for 5 minutes at 4°C. The resulting pellet was gently dissociated using repeated pipetting to generate a single-cell suspension in TrypLE Express dissociation medium (Gibco) using a pipette tip. The suspension was incubated at 37°C for 5 minutes to facilitate dissociation. The reaction was then halted by adding DMEM with 10% (v/v) FBS, and the suspension was centrifuged at 300 g for 5 minutes at 4°C. After discarding the supernatant, the pellet was resuspended in fresh Matrigel in a 24-well plate. Each well contained six drops of Matrigel containing crypts (about 50 µl per well), which were then inverted to polymerise at 37°C in a CO_2_ incubator. After polymerization, 500 µL of Expansion Media (EM) was added to the culture. For differentiation, PM was applied for 7 days at passage P3 and P6, followed by DM for an additional 7 days. The organoid culture was maintained up to P12 in EM. Table 1 describes the media composition; the colon and duodenum shared the same EM, with only the PM and DM deviating between tissues. The basal media used is the same as detailed in previous articles: Advanced DMEM/F12 with 1% (v/v) Hepes, 1% (v/v) Glutamax and 1% (v/v) penicillin-streptomycin and 0.25 µg/ml amphotericin-B (15,19,20).

**Table 1.**
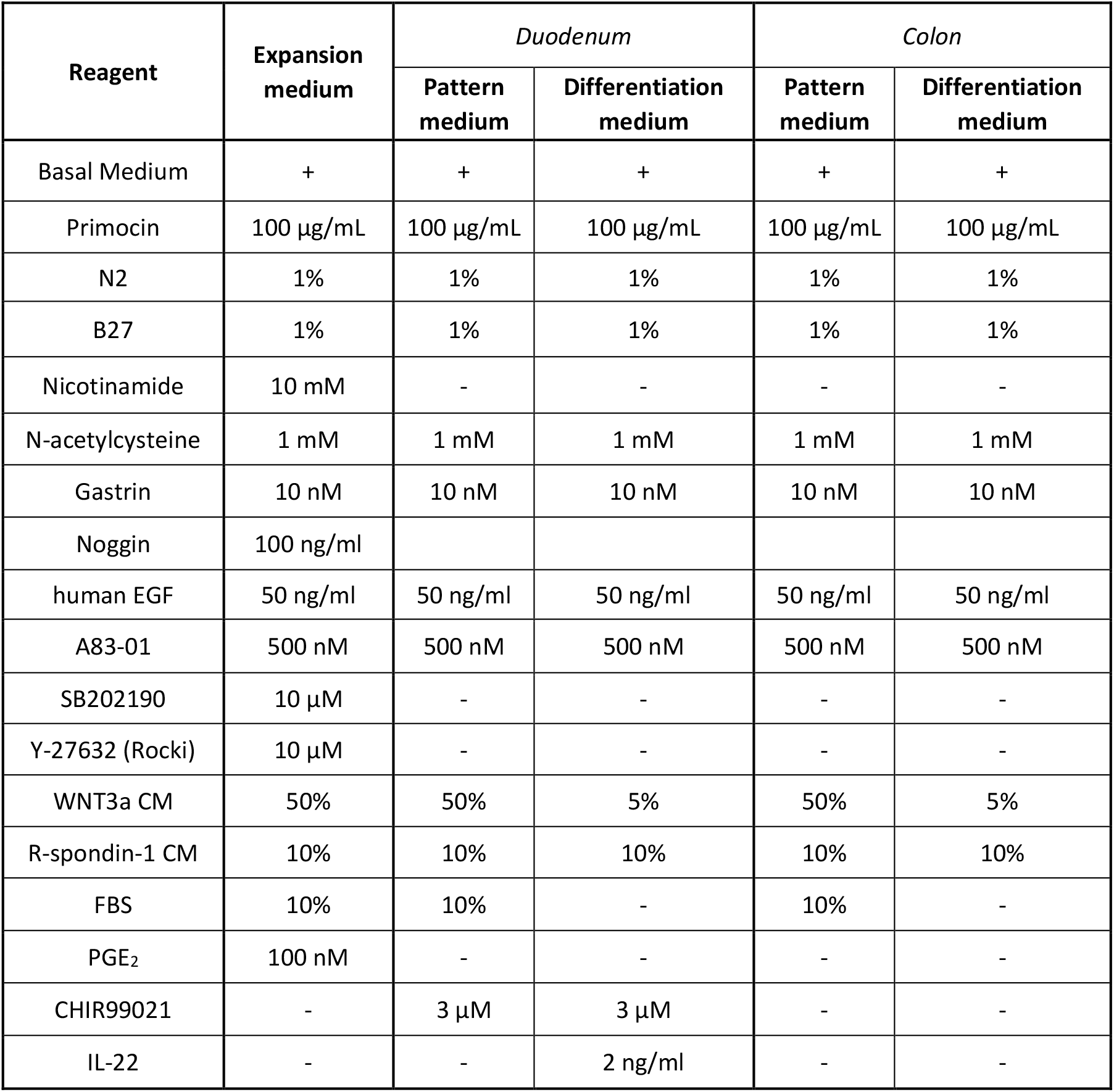
Expansion, Pattern and Differentiation Medium of canine intestinal organoids (duodenum and colon)

| Reagent | Expansion medium | *Duodenum* |  | *Colon* |  |
| --- | --- | --- | --- | --- | --- |
|  |  | Pattern medium | Differentiation medium | Pattern medium | Differentiation medium |
| Basal Medium | + | + | + | + | + |
| Primocin | 100 µg/mL | 100 µg/mL | 100 µg/mL | 100 µg/mL | 100 µg/mL |
| N2 | 1% | 1% | 1% | 1% | 1% |
| B27 | 1% | 1% | 1% | 1% | 1% |
| Nicotinamide | 10 mM | - | - | - | - |
| N-acetylcysteine | 1 mM | 1 mM | 1 mM | 1 mM | 1 mM |
| Gastrin | 10 nM | 10 nM | 10 nM | 10 nM | 10 nM |
| Noggin | 100 ng/ml |  |  |  |  |
| human EGF | 50 ng/ml | 50 ng/ml | 50 ng/ml | 50 ng/ml | 50 ng/ml |
| A83-01 | 500 nM | 500 nM | 500 nM | 500 nM | 500 nM |
| SB202190 | 10 µM | - | - | - | - |
| Y-27632 (Rocki) | 10 µM | - | - | - | - |
| WNT3a CM | 50% | 50% | 5% | 50% | 5% |
| R-spondin-1 CM | 10% | 10% | 10% | 10% | 10% |
| FBS | 10% | 10% | - | 10% | - |
| PGE<sub>2</sub> | 100 nM | - | - | - | - |
| CHIR99021 | - | 3 µM | 3 µM | - | - |
| IL-22 | - | - | 2 ng/ml | - | - |

For cryopreservation, recovery cell freezing media (Invitrogen) was used to aid in the recovery of organoids post-thaw. Approximately 1 million organoids were frozen per vial and stored in liquid nitrogen.

### 2.3. RNA Extraction and Quantitative Real-Time Polymerase Chain Reaction Analysis

Total RNA was extracted from organoids and tissue using the RNeasy Micro- and Mini-Kit (QIAGEN) respectively, in accordance with the manufacturer’s recommendations. The cell pellet was initially snap-frozen and subsequently resuspended in 350 µL of RLT lysis buffer containing 1% (v/v) β-mercaptoethanol. For tissue samples, the purified RNA was resuspended in 30 μL of RNase-free water, while for organoids, 50 μL was used. Complementary DNA (cDNA) was synthesized from total RNA using the iScript™ cDNA Synthesis Kit (Bio-Rad), according to the manufacturer’s instructions. The cDNA was subsequently employed in qPCR analysis using iTaq™ Universal SYBR® Green Supermix (Bio-Rad). The samples were analyzed on Bio-Rad CFX384 Touch Real-Time PCR systems (Bio-Rad) with Bio-Rad CFX Manager 1.1. Gene expression of the target markers was normalized to the mean of three reference genes; Ribosomal Protein L13 (RPL13), Ribosomal Protein S19 (RPS19), and Beta-2-Microglobulin (B2M).

The expression of five key intestinal markers was analysed; Villin 1 (VIL1), a protein component of the brush border of intestinal epithelial cells; Sucrase-isomaltase (SI), an enzyme located in the brush border membrane involved in the digestion of sucrose and starch; Leucine-rich repeat-containing G protein-coupled receptor 5 (LGR5), a marker of intestinal stem cells; Chromogranin A (CHGA), a protein found in enteroendocrine cells; and Mucin 2 (MUC2), a gel-forming mucin that acts as a physical and chemical barrier produced by Goblet cells, protecting the underlying epithelial cells. The forward and reverse primer sequences and Gene IDs for all analysed genes, including the target markers and the reference genes, are listed in Table 2.

**Table 2.**
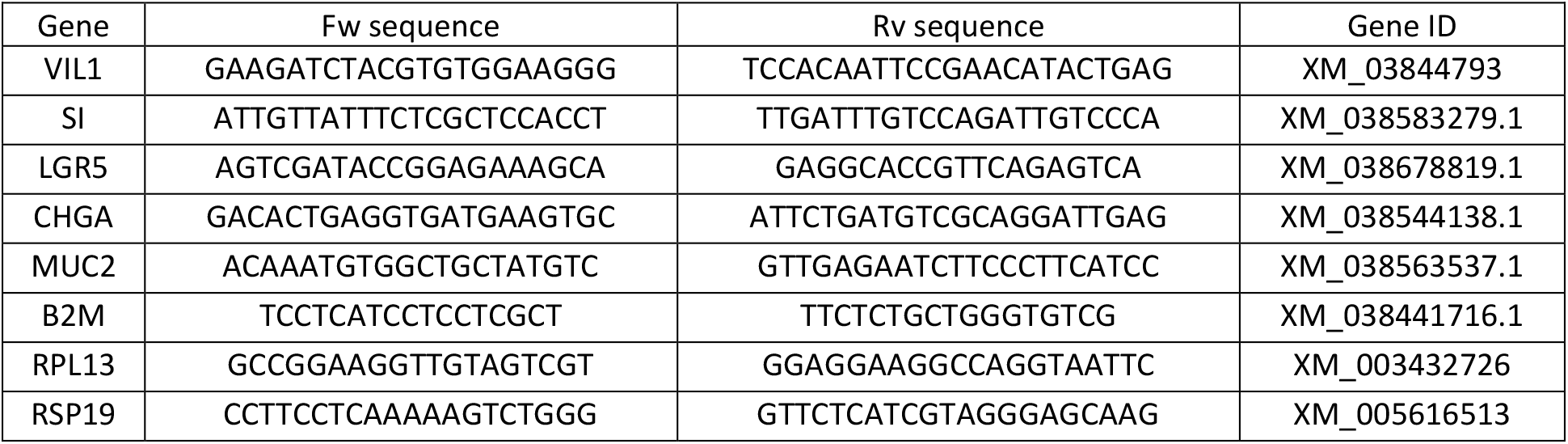
Primer sequences and Gene IDs of the intestinal markers (VIL1, SI, LGR5, CHGA, MUC2) and the reference genes (B2M, RPL13, RSP19) used for quantitative real-time PCR analysis.

### 2.4. Immunohistochemistry staining

Organoids embedded in Matrigel were carefully harvested from 24-well plates using Cell Recovery Solution (CRS) (Corning®) and transferred to 15 mL Falcon tubes. The tubes containing the organoids and CRS were incubated on a tube roller at 4°C for 30 minutes. Following incubation, the tubes were filled with cold PBS and centrifuged at 50 g for 3 minutes at 4ºC. The supernatant was then removed, and the organoids were fixed in 4% (v/v) paraformaldehyde for 45 minutes at 4°C. After fixation, the organoids were washed with 70% (v/v) ethanol to complete the preparation process.

For histological analysis, organoids were embedded in paraffin blocks, sectioned, adhered to slides in a 55°C oven and dehydrated. Antigen retrieval was performed by heating the slides in TE buffer (pH 9.0) until boiling, followed by rinsing in PBS at room temperature. Permeabilization was carried out in PBS containing 0.1% (v/v) Triton X-100 (PBST) for 10 minutes, after which the slides were blocked with 10% (v/v) normal goat serum for 30 minutes. The samples were incubated with primary antibodies listed in Table 3 overnight at 4°C in 10% (v/v) normal goat serum. The slides were then washed with PBST, followed by incubation with secondary antibodies (568 Goat anti-rabbit and 568 Goat anti-mouse) for 1 hour in the dark at room temperature. After additional PBS washes, the slides were stained with DAPI for 10 minutes, rinsed again with PBS, and finally mounted for imaging. Slides were imaged using the Leica DM4000 microscope.

**Table 3.**
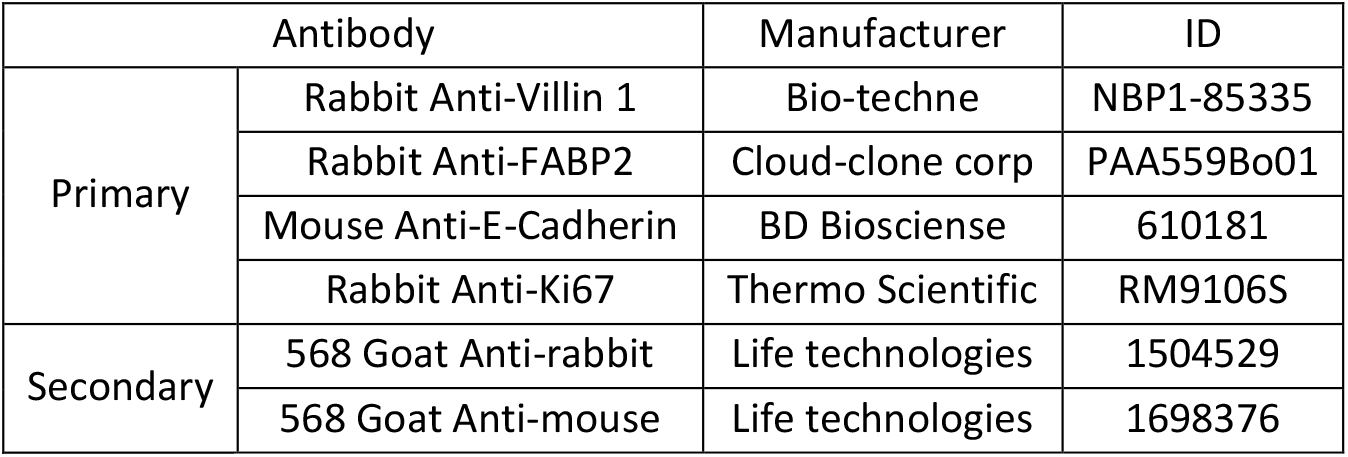
Primary and secondary antibodies used for immunohistochemistry.

### 2.5. CFTR functional assay using canine enteroids

Enteroids and colonoids were passaged and seeded in Matrigel into 24-well plates as previously described. At P5 and P10, the organoids were transferred to 96-well plates (one drop per well) to study four conditions: 2.5 µM, 5 µM, and 10 µM forskolin, and a matched vehicle control (DMSO) at an equivalent final dilution. To each well, 100 µL of the forskolin solution at the respective concentration or DMSO for the control were added. Representative images of enteroids were captured after 0, 1, 2, and 3 hours using a Life Technologies EVOS™ FL microscope.

### 2.6. Statistical Evaluation

Statistical analyses were performed in IBM SPSS Statistics (v30). Because biological replicates were limited (n = 2 donors per group) and measurements were acquired as technical duplicates, we treated donors as the experimental unit and summarized technical replicates at the donor level. Normality was assessed using the Shapiro–Wilk test. Given the small sample size and the presence of non-normal distributions, non-parametric tests were selected for all comparisons. For three-group comparisons (EM, PM, DM), the Kruskal–Wallis test was used. Where appropriate, pairwise post-hoc comparisons were performed with the Mann–Whitney U test with Bonferroni correction. For paired comparisons between P3 and P6 within the same donor, we applied the Wilcoxon signed-rank test.

## 3. Results

### 3.1. PGE_2_ Promotes Canine Intestinal Organoid Growth in a Dose-Dependent Manner

Canine intestinal organoids derived from duodenal and colonic tissues were successfully generated and cultured in EM, as previously described. The composition of the EM was similar to that used in prior studies on dogs (15,19,20), with the addition of PGE_2_. Organoids cultured with higher concentrations of PGE_2_ (100 nM compared to 10 nM) demonstrated improved growth and greater proliferation. Consequently, a concentration of 100 nM PGE_2_ was maintained in the EM for both duodenal and colonic organoids. Figure 2 illustrates the morphology and growth of duodenal and colonic organoids cultured without PGE_2_ and with 10 nM and 100 nM of PGE_2_.

**Figure 2.**
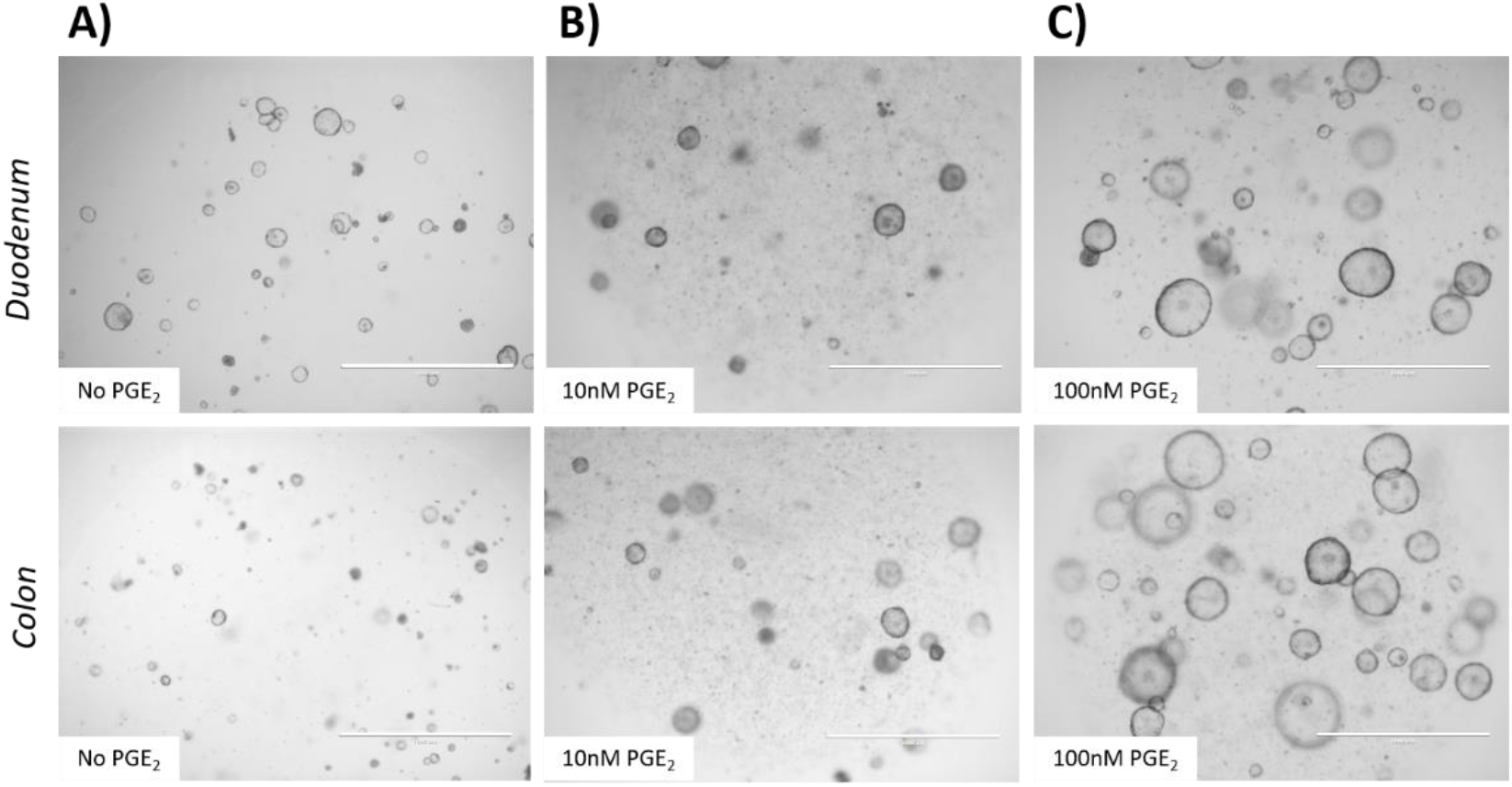
Representative images of canine intestinal organoids cultured with different PGE2 concentrations. Images were acquired at passage 1 (P1) on day 3 of culture. (A) Duodenum (top) and colon (bottom) cultured without PGE_2_. (B) Duodenum (top) and colon (bottom) cultured with 10 nM PGE_2_. (C) Duodenum (top) and colon (bottom) cultured with 100 nM PGE_2_. Scale bar = 1,000 µm

### 3.2. Expansion Medium Enables Proliferation and Growth of Canine Intestinal Organoids, While Pattern and Differentiation Media Promote Their Differentiation

The organoids were maintained in culture up to P12, with a split ratio of 1:4 every 7 days, after which they began to degenerate. Assuming a constant expansion of 4-fold per passage, this corresponds to a cumulative expansion potential of approximately 1.7 × 10^7^-fold by P12, illustrating the high yield achievable with this system. At P3 and P6, differentiation of duodenal and colonic organoids was initiated in two phases: first using PM for 7 days, followed by DM for an additional 7 days. In contrast to the colon, CHIR99021 and IL-22 were included in the DM for the duodenum, as described by He *et al*. (2022) and Almeqdadi *et al*. (2019) (21,26). CHIR99021, a GSK-3 inhibitor that activates the Wnt/β-catenin pathway, was added to maintain stem-cell signalling and promote crypt-like budding characteristic of the small-intestinal epithelium, while IL-22 was added to support Paneth-cell differentiation. During differentiation, the organoids became denser, developed an irregular surface, and acquired a darker appearance, particularly between days 4 and 7 of the differentiation process. Figure 3 shows representative duodenal and colonic organoids cultured in EM, PM, and DM.

**Figure 3.**
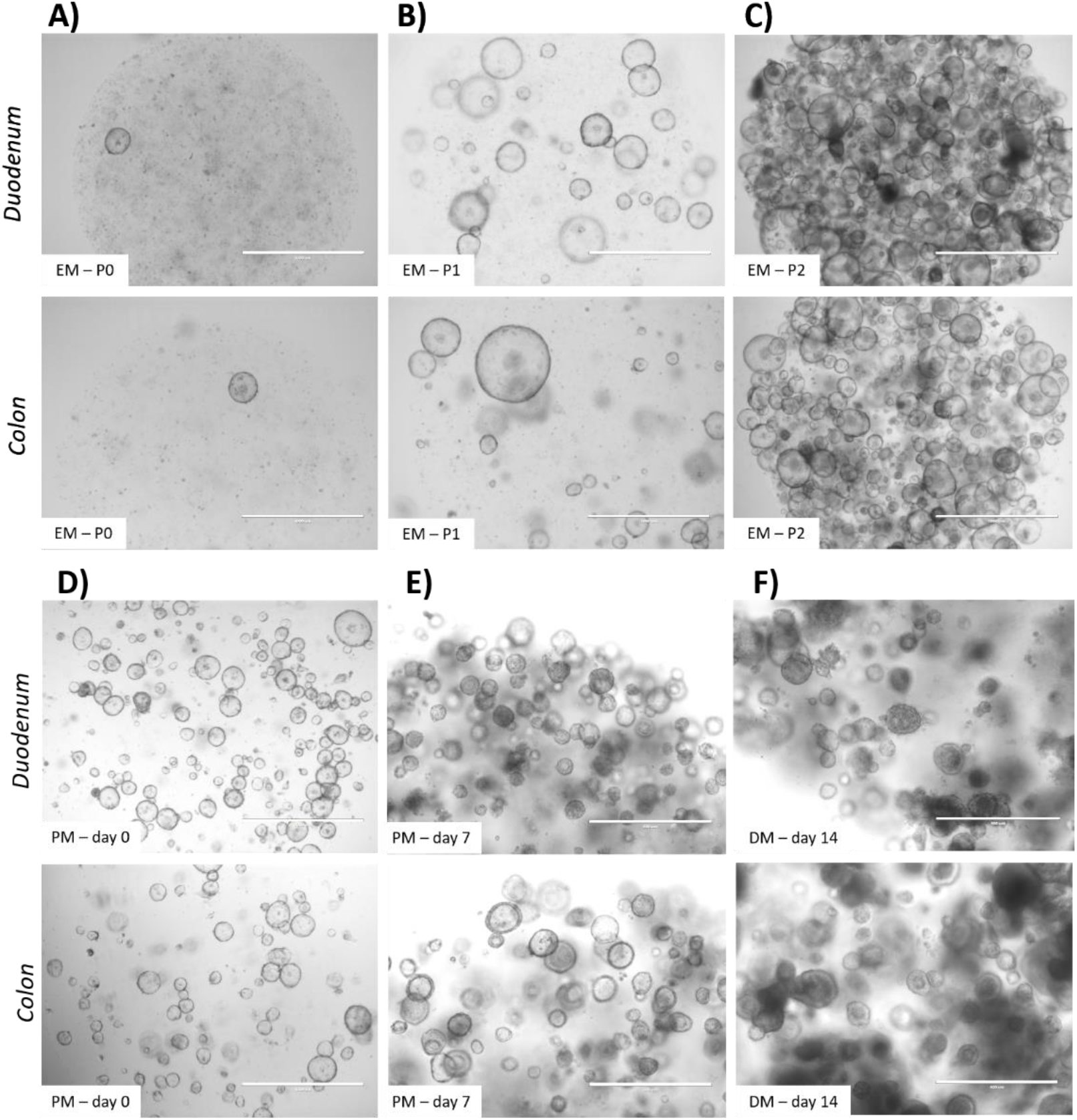
Representative images of duodenal and colonic organoids. cultured in expansion medium (EM) at P0 (A), P1 (B), and P2 (C). Representative images of duodenal and colonic organoids cultured in pattern medium (PM) at day 0 (D), PM at day 7 (E), and differentiation medium (DM) at day 14—seven days after exposure to differentiation medium (F). Scale bar = 1,000 µm

To evaluate the impact of media composition and passage number on the cellular composition of organoids, qRT-PCR analyses were performed for VIL1, SI, LGR5, CHGA, and MUC2. Non-parametric analyses showed no statistically significant differences across media (EM, PM, DM) or between passages (P3 vs. P6) in either tissue (all p > 0.05; Kruskal–Wallis and Wilcoxon tests), which is consistent with the small sample size (n = 2 donors) and inter-donor variability. Nevertheless, consistent biological trends were evident (Figure 4), duodenum-derived organoids generally exhibited higher expression levels of all markers compared to colon-derived organoids. In duodenal organoids, most markers, including absorptive (VIL1, SI), secretory (CHGA, MUC2), and the stem cell marker LGR5, showed higher expression under PM, suggesting activation of epithelial differentiation programs together with maintenance of a stem/progenitor population. In contrast, colon-derived organoids displayed a different pattern, with absorptive markers (VIL1 and SI) and the enteroendocrine marker CHGA reaching higher expression under DM, while LGR5 and MUC2 remained more elevated under PM. These findings suggest segment-specific responses to culture conditions between small intestinal and colonic organoids. Overall, there was a tendency towards higher marker expression in PM and/or DM compared to EM in both tissue types. These patterns may reflect region-specific epithelial identities that are retained in intestinal organoid cultures, as organoids preserve key characteristics of their tissue of origin (3).

**Figure 4.**
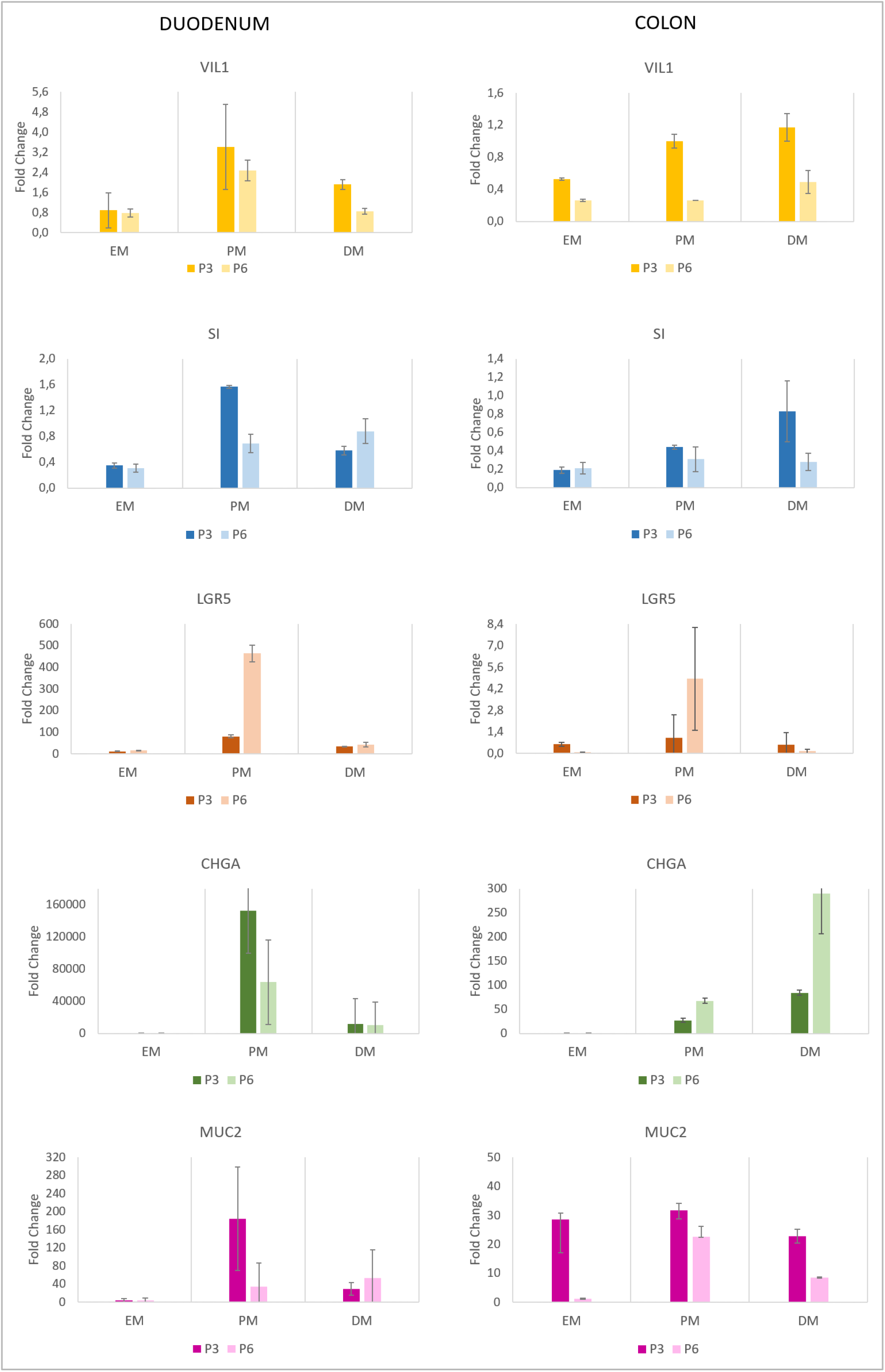
Relative expression of SI (sucrase-isomaltase), VIL1 (villin 1), CHGA (chromogranin A), LGR5 (leucine-rich repeat-containing G protein-coupled receptor 5), and MUC2 (mucin 2) in organoids from duodenum (left) and colon (right) at passage 3 (P3) and passage 6 (P6). Gene expression was normalized to native tissue from the corresponding intestinal segment. Statistical significance for paired comparisons between P3 and P6 was determined using the Wilcoxon signed-rank test. The Kruskal-Wallis test was used in the overall statistical analysis as described in the Methods section. One biological sample with two technical replicates, values are presented as mean ± SD for each intestinal segment (two dogs).

To further investigate the protein expression and localization of specific intestinal markers under different culture conditions, we performed immunohistochemistry on control tissue, organoids cultured in EM, and organoids cultured in DM. Figure 5 displays confocal single-plane images of duodenum tissue and organoids stained for VIL1, FABP2, E-cadherin, and the proliferation marker Ki-67 (red). Similarly, Figure 6 presents confocal single-plane images of colon tissue and organoids stained for the same markers under the same culture conditions.

**Figure 5.**
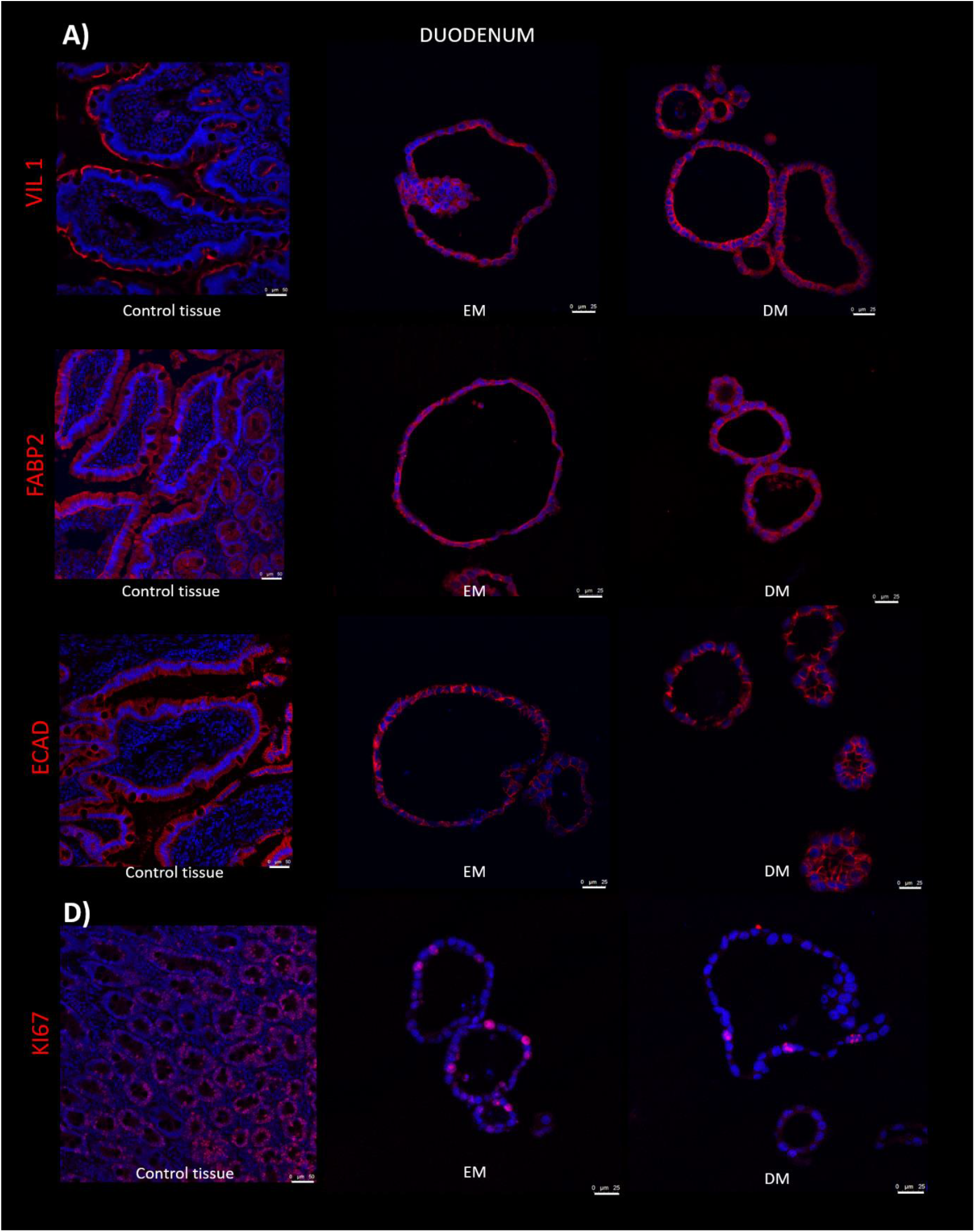
Duodenum confocal single-plane image showing intestinal markers villin (A), FABP2 (B), E-cadherin (D) and KI-67 (D) (red) in control tissue, expansion medium (EM) and differentiation medium (DM). Scale bar = 50 µm for tissue and 25 µm for organoids.

**Figure 6.**
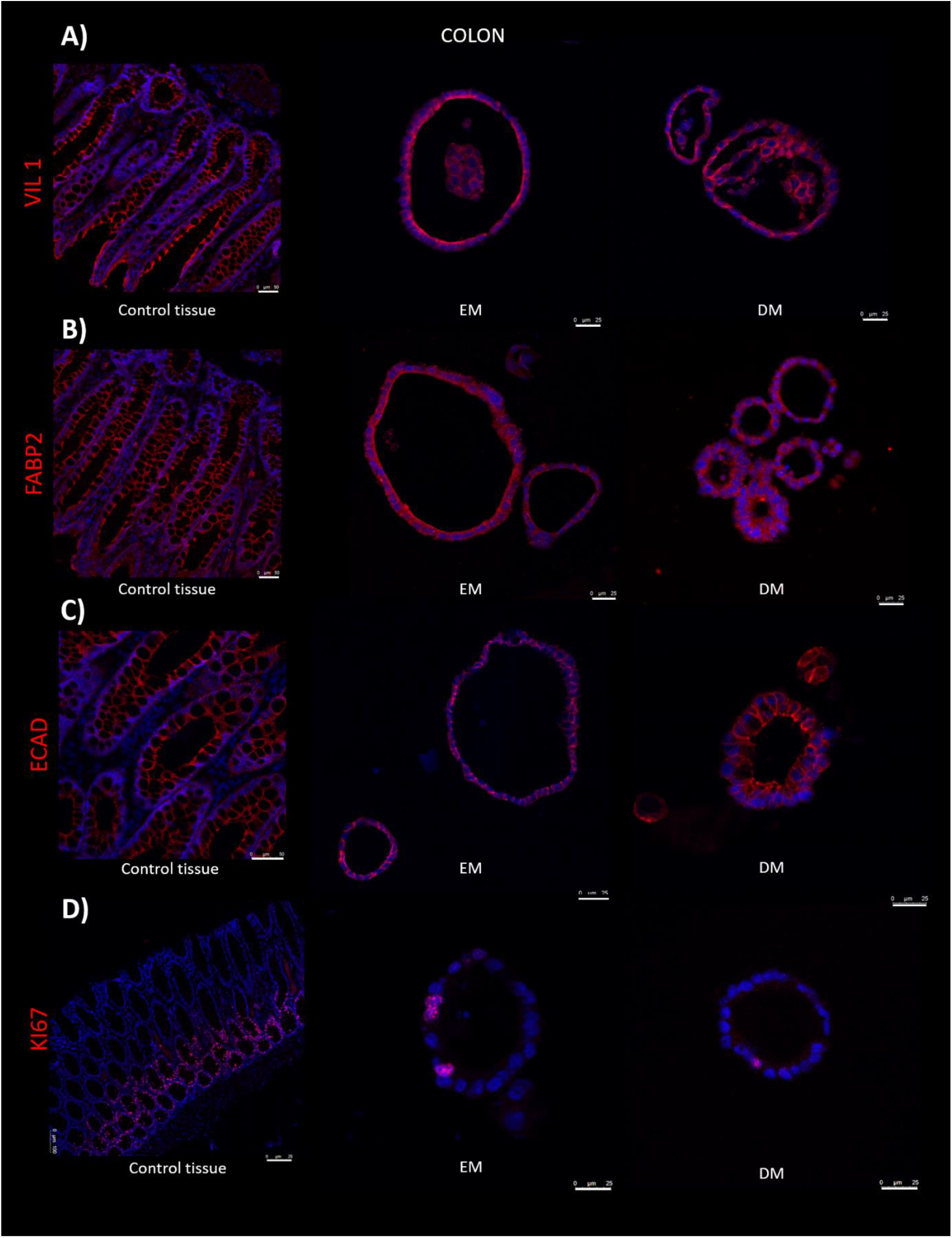
Colon confocal single-plane image showing intestinal markers villin (A), FABP2 (B), E-cadherin (D) and KI-67 (D) (red) in control tissue, expansion medium (EM) and differentiation medium (DM). Scale bar = 50 µm for tissue and 25 µm for organoids.

Qualitative assessment of the immunohistochemistry images was performed by visual inspection using fluorescence microscopy. Organoids cultured in EM exhibited expression of all examined markers (VIL1 and FABP2, markers of absorptive enterocyte differentiation; E-cadherin, indicative of epithelial organization; and the proliferation marker Ki-67), similar to those observed in DM. The presence of VIL1 and FABP2 in both conditions indicates the establishment of an absorptive epithelial phenotype, while E-cadherin expression confirms maintenance of epithelial integrity under both culture conditions. Notably, the proliferation marker Ki-67 appeared qualitatively reduced in DM-cultured organoids compared to EM in both duodenal and colonic samples.

### 3.3. Forskolin-Induced Swelling Demonstrates Functional CFTR Activity in Canine Intestinal Organoids in a Dose-Dependent Manner

To assess organoid functionality, we conducted a swelling assay using forskolin, a cAMP agonist that activates CFTR chloride channels in intestinal epithelial cells, causing canine enteroids to swell (15,27). Since CFTR activation relies on intracellular cAMP levels, forskolin promotes water flow into the enteroid lumen, leading to swelling. This makes the assay an indirect way to measure CFTR function in enteroids. The swelling was observed up to three hours, increasing over time, confirming that CFTR in the canine enteroids is functional, as seen in intact tissue. Figure 7 illustrates the evolution of intestinal organoids with forskolin at doses of 2.5, 5, and 10 µL, along with the DMSO control, over a period of 3 hours.

**Figure 7.**
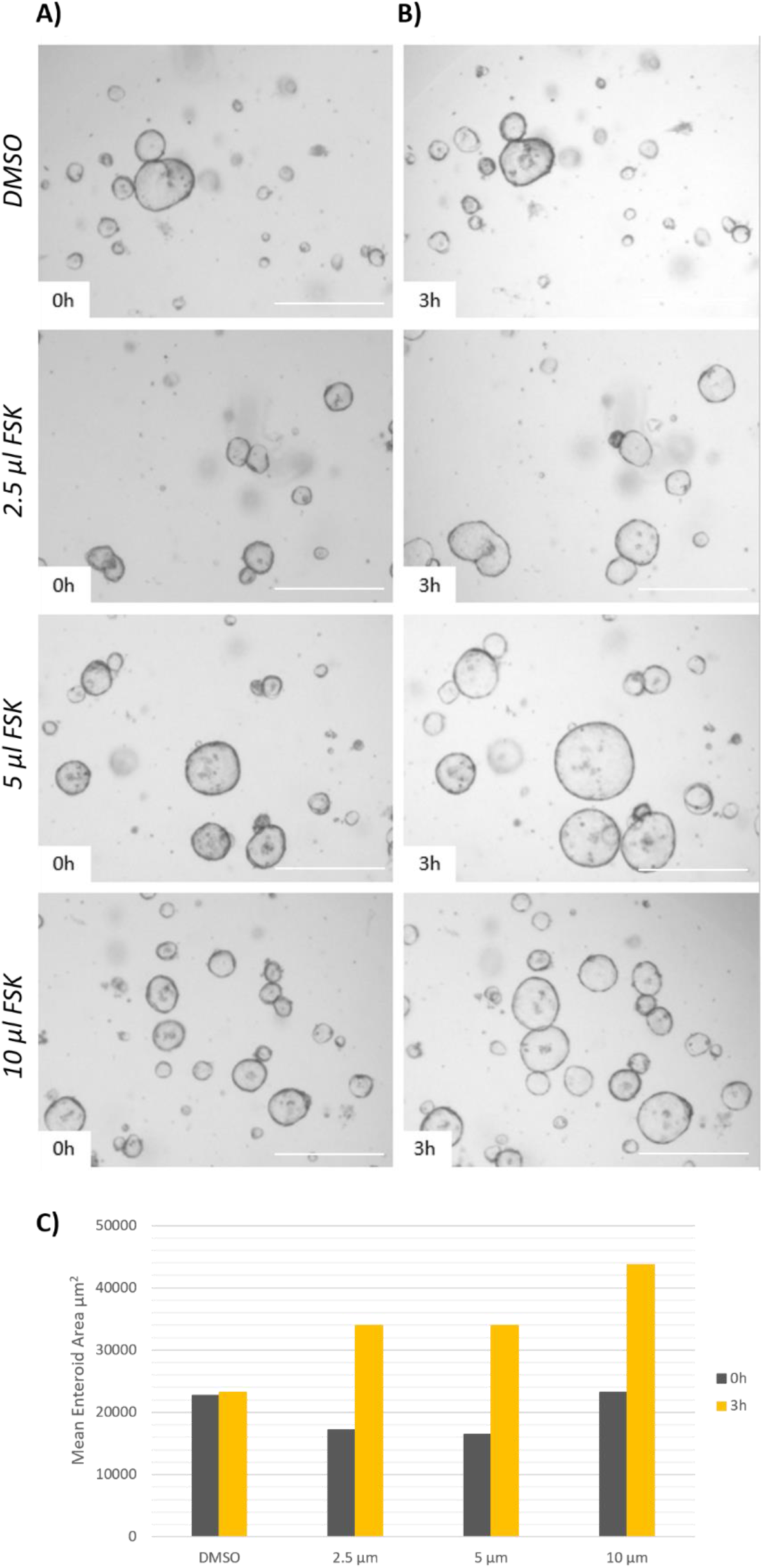
Forskolin induces time-dependent swelling of canine enteroids, suggesting the presence of functional CFTR. Representative images of enteroids were captured at baseline (A) and 3 hours after forskolin was introduced into the medium (B). A graph showing the mean area of enteroids, measured from 10 organoids per field across *n* = 5 fields per condition, is presented (C). Scale bar = 1,000 µm, 4x magnification

## 4. Discussion

This study successfully established a protocol for the isolation, long-term culture, and differentiation of canine intestinal organoids derived from both the duodenum and colon. The use of an optimized growth factor cocktail, as previously described (7,8), in conjunction with a three-dimensional Matrigel matrix, facilitated the formation of self-renewing organoid structures that recapitulate key aspects of the native canine intestinal epithelium. Notably, our findings indicate that the inclusion of PGE_2_ (100 nM) in the EM significantly promoted organoid growth compared to conditions without PGE_2_ or with a lower concentration (10 nM). The enhanced growth observed with 100 nM PGE_2_ suggests that this concentration provides an optimal signalling environment for the expansion of canine intestinal stem and progenitor cells *in vitro*. This finding is likely mediated through PGE_2_-EP4-dependent activation of β-catenin signalling, which enhances stem cell proliferation and epithelial budding in intestinal models (24).

Morphological and functional characterization further validated the utility of our canine organoid model. While statistical analysis of gene expression across different culture media (EM, PM, DM) and passages (P3, P6) did not reveal significant differences in the expression of selected epithelial markers (VIL1, SI, CHGA, LGR5, MUC2), several biologically relevant trends were observed. Because duodenal and colonic organoids were analysed relative to their own tissue references, no direct comparison between tissues was performed. In duodenum-derived organoids, all examined markers (VIL1, SI, LGR5, CHGA, MUC2) were more highly expressed in the PM compared to the DM. This could suggest that the removal of certain components present in EM—such as nicotinamide, Noggin, SB202190, Y-27632 and PGE_2_—may play a more decisive role in promoting differentiation than the addition of CHIR99021 and IL-22 (28,29), which are unique to the DM. IL-22 has been associated with Paneth cell development in human intestinal organoids (21); however, Paneth cell markers were not evaluated in the present study. In dogs, the existence of classical Paneth cells remains uncertain, and further investigation of Paneth or Paneth-like cell differentiation in canine organoids is warranted. The addition of CHIR99021 may have prolonged Wnt-driven stem-cell signalling during differentiation. However, given the limited sample size, these observations should be interpreted with caution. In contrast, colon-derived organoids showed a distinct pattern of response: DM induced higher expression of absorptive enterocyte markers (VIL1 and SI), as well as secretory lineage markers, including CHGA, whereas MUC2 expression was higher in PM than in DM, suggesting preferential support of goblet cell–associated programs under PM conditions. Although SI is primarily a small-intestinal enterocyte marker and not normally expressed in the adult colon (30), its detection in colonic organoids likely reflects culture-induced activation of absorptive enterocyte gene programs. Notably, LGR5 expression remained elevated in PM, consistent with the known dependence of LGR5^+^ intestinal stem cells on active Wnt signalling, which is present in EM and PM but absent in DM. Together, these findings underscore the region-specific sensitivity of canine intestinal organoids to media composition, emphasizing the need to tailor differentiation protocols to the tissue of origin. These transcriptional trends were supported by immunohistochemistry, which confirmed epithelial identity and enterocyte differentiation through the detection of Villin-1, FABP2 and E-cadherin, and revealed reduced Ki-67 staining under DM conditions, indicative of decreased proliferation and progression toward a more differentiated epithelial state.

The absence of statistically significant changes in marker expression across passages (P3 and P6) suggests that the established culture conditions support a relatively stable cellular phenotype over short-term passaging. However, a significant limitation of this study is the small sample size, with organoids derived from only two individual dogs (one duodenum and one colon). This limited sample size inherently reduces the statistical power of our analyses, making it challenging to draw definitive conclusions about the effects of different culture conditions and passages on marker expression. The observed biological trends, while intriguing, require validation in a larger cohort to determine their consistency and statistical significance.

Furthermore, our functional assessment using a forskolin-induced swelling assay demonstrated the presence of functional CFTR chloride channels in both duodenal and colonic canine intestinal organoids. The observed time and dose-dependent increase in organoid swelling upon forskolin stimulation confirms that these *in vitro* models retain a key physiological function characteristic of the native intestinal epithelium. In addition, we have successfully cryopreserved the organoids derived from both the duodenum and colon, ensuring a valuable resource for ongoing and future investigations.

## 5. Conclusion

This study establishes a protocol for the generation and characterization of canine intestinal organoids from both the duodenum and colon. Functional validation through CFTR activity and detection of key epithelial markers demonstrates the presence of major intestinal cell types within the cultures, supporting the physiological relevance of this model for *in vitro* studies. However, as these markers were also detected under expansion conditions, further work is needed to confirm full functional differentiation and to refine lineage-specific maturation protocols. Notably, the addition of PGE_2_ (100 nM) significantly enhanced organoid proliferation in expansion conditions, while reduction of Wnt signalling and removal of nicotinamide, Noggin, SB202190, Y-27632, and PGE_2_ promoted differentiation in duodenal and colonic cultures. Although the limited sample size constrains statistical power, the observed trends provide a foundation for future studies. Further optimization of differentiation timing may enhance marker expression and refine culture protocols.

